# Double-targeted knockdown of miR-21 and CXCR4 inhibits malignant glioma progression by suppression of the PI3K/AKT and Raf/MEK/ERK pathways

**DOI:** 10.1101/2020.04.14.040568

**Authors:** Feijiao Liu, Bo Yang

## Abstract

**Background:** Currently, miR-21 and CXCR4 are being extensively investigated as two unrelated key regulators in glioma malignancy. In this study, we investigated the combined effect of these two factors on glioma progression.

**Methods:** We confirmed the expression of miR-21 and CXCR4 in malignant glioma tissue and glioma cells with qRT-PCR and western blotting. Single-targeted knockdown of miR-21 and CXCR4, as well as double-targeted knockdown of miR-21 and CXCR4 lentiviral vectors were constructed and they were transfected to U87 and U251 cells. Cell proliferation, apoptosis, invasion, and migration from different treatment groups were assessed by MTT assay, Flow Cytometry analysis, Transwell analysis, and Scratch assay, respectively. U87 xenograft mice were constructed to detect roles and potential mechanisms of miR-21 and CXCR4 in malignant glioma tumor growth.

**Results:** The expression of miR-21 and CXCR4 was increased in tumor tissues and cell lines. Inhibition of miR-21, CXCR4, and miR-21 and CXCR4 together all reduced the migration, invasiveness, proliferation and enhanced apoptosis in glioma cells, as well as reduced tumor volume and mass in xenograft model. The inhibition effect was strongest in double-targeted knockdown of miR-21 and CXCR4 group, whose downstream pathways involved in AKT axis and ERK axis activation.

**Conclusions:** Our findings reported that double-targeted knockdown of miR-21 and CXCR4 could more effectively inhibit the proliferation, migration, invasion and growth of transplanted tumor and promote cell apoptosis, which were involved in the PI3K/AKT and Raf/MEK/ERK signaling pathways.

## Introduction

Glioma is the most common and feared type of tumor, with an annual incidence of 1 per 10,000 population (Yang et al., 2018). Currently, treatments for malignant glioma are mainly surgical resection, followed by chemotherapy or radiotherapy (Andrea et al., 2013). However, these comprehensive treatments are not very effective and the median survival of patients afflicted with malignant glioma is less than 1 year, due to the aggressive proliferation and insidious invasion of cells (Skalsky and Cullen 2011, Põlajeva et al., 2012). Thus, recent studies are focusing on the factors that can effectively control the invasive and proliferative capacity of malignant glioma cells to develop targeted salvage treatments to improve patients’ life quality.

miR-21 is an oncomiR which attracts many attentions, because it takes part in almost all steps of tumor growth and metastasis [6]. Evidence shows that miR-21 is overexpressed in gliomas and glioma cells compared to normal tissues, and its expression level is positively correlated with glioma grade (Tao et al., 2013, Lei et al., 2014). Elevated miR-21 expression leads to increased glioma cell proliferation invasion, and also chemo-resistance, which indicates poor prognosis and tumor recurrence of glioma patients (Gabriely et al., 2008, Kwak et al., 2011, Melnik 2015, Hermansen et al., 2016), On the other hand, downregulation of miR-21 leads to repression of anti-apoptotic capacity, reduction of migratory and invasion, as well as increase of chemical-induced death in glioma cells (Gabriely *et al.* 2008, Loges et al., 2012, Luo et al., 2017, Shao et al., 2017). All of these make miR-21 not only a potential glioma marker for diagnosis and prognosis, but a target for novel therapeutic intervention.

Another factor that draws great attention for playing a crucial role in malignant glioma biological regulation is C-X-C Chemokine Receptor 4 (CXCR4), a transmembrane G-protein-coupled receptor. Statistics shows that CXCR4 level is elevated in malignant glioma when compared to normal cells derived from the same tumor (Zhang et al., 2007). Particularly, its overexpression is concentrated in invading regions, and associated with increased glioma tumor grade and malignancy (Chatterjee et al., 2014, Yang *et al.* 2018). Numerous studies suggest that the overexpression of CXCR4 facilitates proliferation, angiogenesis, invasion, metastasis, as well as chemotherapy and radiotherapy resistance of glioma in several glioma cell lines and mouse models (Rubin et al., 2003, Bajetto et al., 2006, Redjal et al., 2006, Mercurio et al., 2016, Gagner et al., 2017, Eckert et al., 2018). Administration of either CXCR4 neutralizing antibody or CXCR4 siRNA impairs the enhanced glioma malignancy (proliferation, angiogenesis, invasion, metastasis, and post-chemotherapy and post-radiotherapy recurrence) and increases median survival in *in vivo* glioma models (Rubin *et al.* 2003, Redjal *et al.* 2006, Esencay et al., 2010, Choi et al., 2014, Mercurio *et al.* 2016, Wang et al., 2016, Yadav et al., 2016, Bathen et al., 2017, Gagner *et al.* 2017, Yang et al., 2017). Thus, the potential of CXCR4 as an anti-tumor target has been a popular research topic.

The two moleculars miR-21 and CXCR4 as a single target respectively have drawn wide attention of researches. However, several studies have shown that in gene therapy of cancers, combination of multigene therapy tends to be more effective than single-gene therapy strategy, which may provide a promising treatment in cancer. Herein, we investigated the combined effects of miR-21 and CXCR4 by lentiviral mediated gene recombinant technology on glioma malignancy. Double-targeted knockdown of miR-21 and CXCR4 was performed in glioma cell lines (U87 and U251) and its effects on glioma malignant progression were evaluated in cells and xenograft mouse models. Knockdown of either miR-21 or CXCR4 decreased glioma proliferation, invasion, and migration and enhanced apoptosis. Double-targeted knockdown generated a significantly enhanced inhibition effect on tumorigenicity even compared to single-targeted knockdown of miR-21 or CXCR4 alone. Also, we found that decrease of tumor progression was related to PI3K/AKT and Raf/MEK/ERK pathways which are responsible for glioma growth, invasiveness, chemo- and radio-therapy resistance, and recurrence (Loges *et al.* 2012, Shao *et al.* 2017). Thus, this study might also pave ways for further research on glioma pathology and physiology.

## Results

### miR-21 and CXCR4 expressions were increased in malignant glioma tissues and cells

In order to determine the effect of double-targeted knockdown of miR-21 and CXCR4 on malignant glioma tumorigenicity, we first confirmed the dysregulation of these two factors in malignant glioma tissues. qRT-PCR was employed to assess the expression of miR-21 and CXCR4 at genetic level. Both miR-21 and CXCR4 average expressions were significantly up-regulated in glioma tissues compared to paracancerous tissues (Fig. 1A-1C). Further, we examined the expression of miR-21 and CXCR4 in two invasive-prone glioma cell lines (U87 and U251) in comparison to a normal glial cell line (HAC). The expression levels of miR-21 and CXCR4 were both significantly increased in U251 and 6-fold higher in U87 (Fig. 1D-1F). The results suggested that miR-21 and CXCR4 might play important roles in mediating malignant glioma aggressiveness, as well as the tumorigenicity.

**Fig.1.**
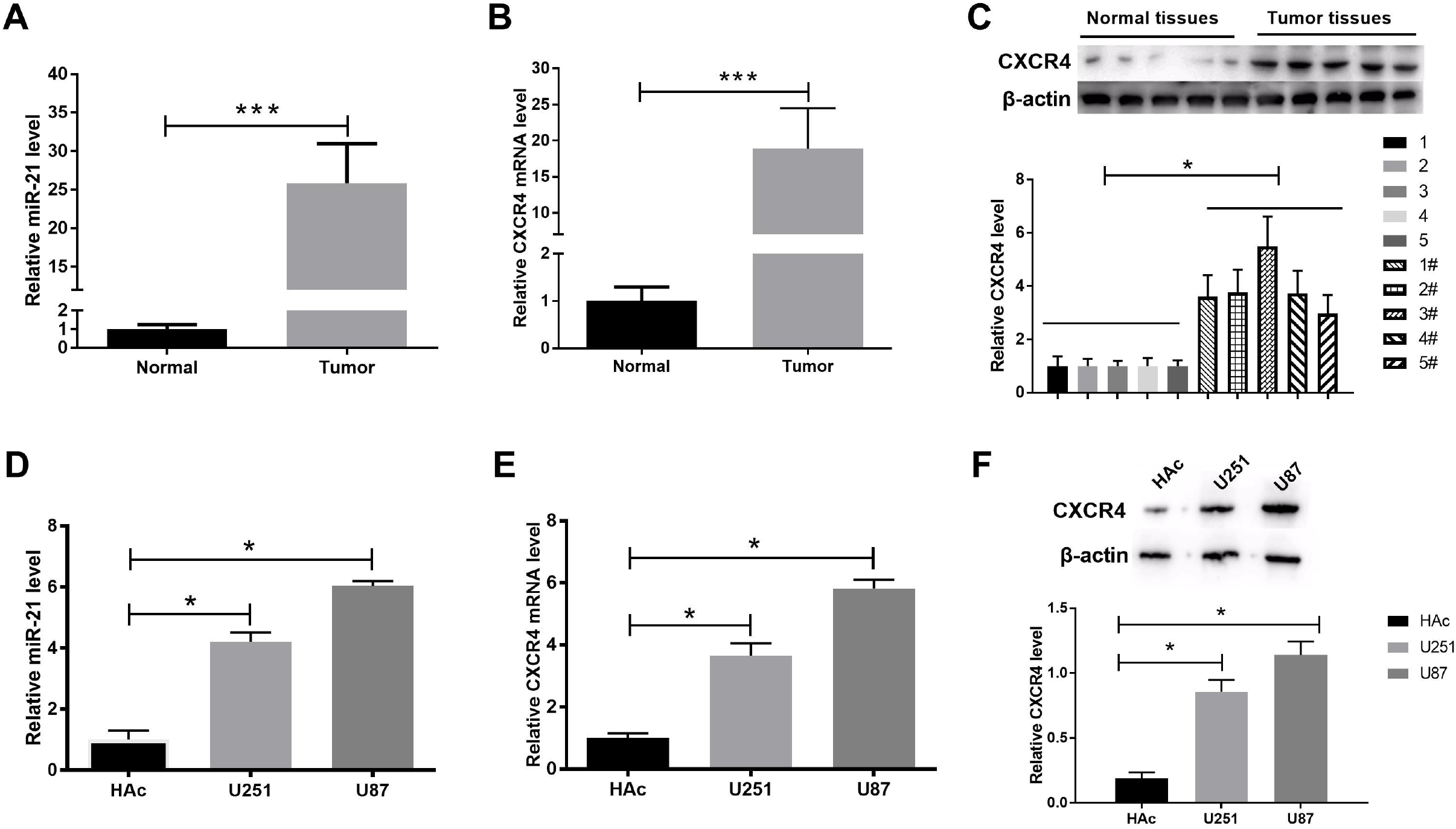
miR-21 and CXCR4 levels in malignant glioma tissue and cells. (**A and B**) The expression of miR-21 and CXCR4 was detected by qRT-PCR in glioma and normal adjacent tissue samples. **(C)** The levels of CXCR4 were determined by western blot in tumor and normal tissue samples. **(D and E)** The expression of miR-21 and CXCR4 was measured by q-RT-PCR in glioma cells (U251 and U87) and control cells (HAC). **(F)** CXCR4 protein levels were tested by western blot. \**P* < 0.05.

### The construction of stable anti-miR-21, sh-CXCR4, and anti-miR-21 + sh-CXCR4 glioma cells

We constructed miR-21 knockdown (anti-miR-21), CXCR4 knockdown (sh-CXCR4), and miR-21 and CXCR4 double-targeted knockdown (anti-miR-21 + sh-CXCR4) U87 and U251 cells to investigate the anti-miR-21 + sh-CXCR4 generated effect on glioma tumorgenicity. qRT-PCR was used to detect the efficiency of miR-21 knockdown at genetic level and data showed that the knockdown efficiency reached more than 70% in both U87 and U251 cells (Fig. 2A). The efficiency of sh-CXCR4 was measured by qRT-PCR and western blot and the results showed that CXCR4 expression level in U251 and U87 cells was significantly repressed compared to negative control and mock group (Fig. 2B-2D). Moreover, the construction of anti-miR-21 + sh-CXCR4 was successful with the efficiency of miR-21 and CXCR4 inhibition in U87 and U251 being comparable to anti-miR-21 or sh-CXCR4 treatment alone generated effects (Fig. 2E-2G).

**Fig.2.**
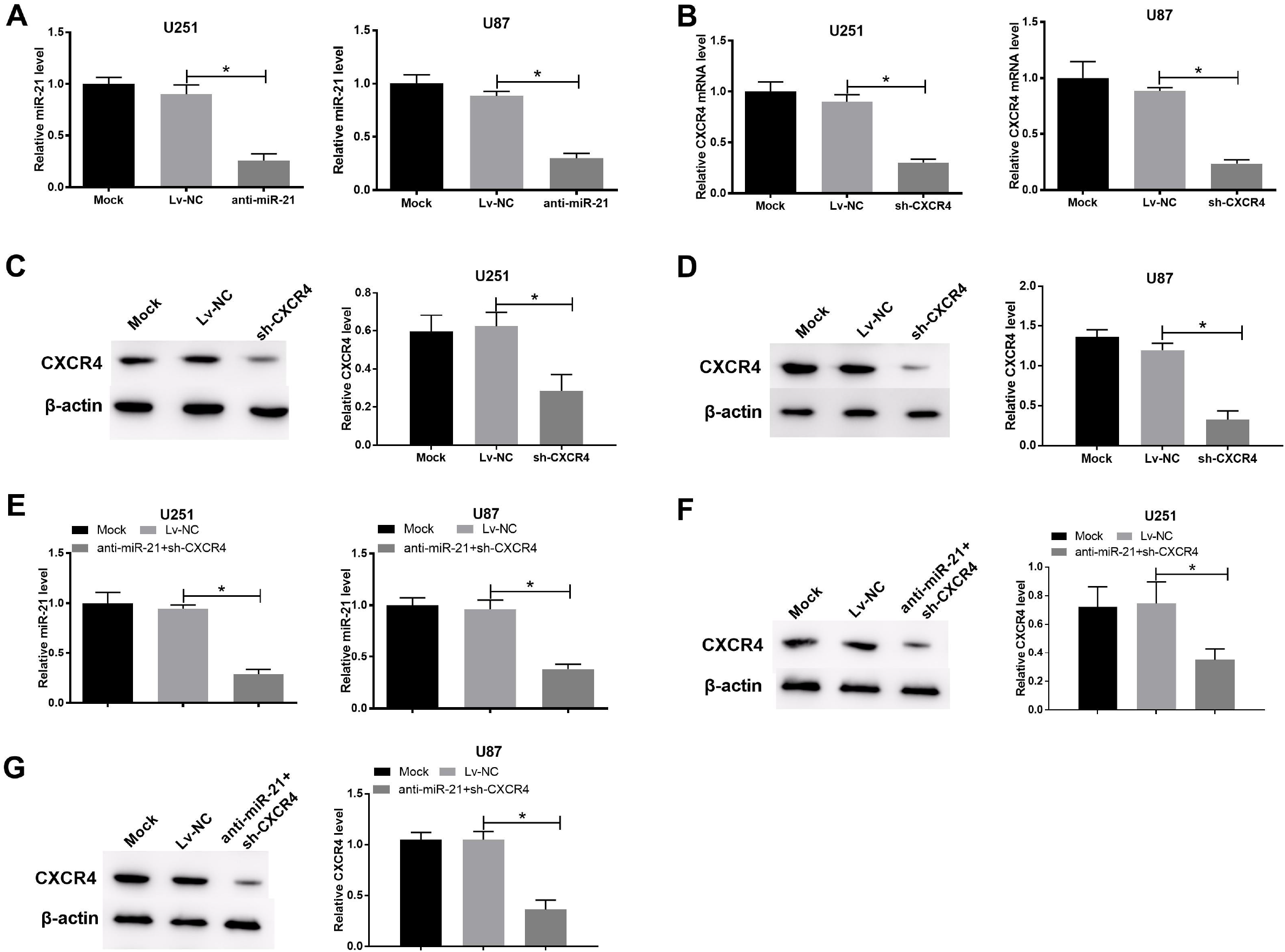
miR-21 and CXCR4 levels of stable glioma cells transfected with anti-mlR-21, sh-CXCR4, or anti-miR-21 + sh-CXCR4. **(A and B)** miR-21 and CXCR4 levels in transfected U251 and U87 cells were measured using qRT-PCR at 72h post-transfection. **(C and D)** CXCR4 protein levels in transfected U251 and U87 cells were measured using western blot. **(E)** miR-21 and CXCR4 levels in transfected U251 and U87 cells were measured using qRT-PCR. **(F and G)** CXCR4 protein levels in transfected U251 and U87 cells were measured using western blot. \**P* < 0.05.

### Double-targeted knockdown of miR-21 and CXCR4 inhibited proliferation and enhanced apoptosis of glioma cells

To explore the roles of miR-21 and CXCR4 in sustaining aggressive proliferation and anti-apoptosis in glioma cells, functional experiments were performed. MTT assay was performed on transfected U87 and U251 cells for the evaluation of cell proliferation at different time points (0, 24, 48, 72 hr). In comparison to negative control group, anti-miR-21 or sh-CXCR4 alone was able to significantly decrease cell proliferation of U87 and U251 (Fig. 3A and Fig. 3B). Interestingly, in anti-miR-21 + sh-CXCR4 group, the inhibition efficiency was significantly higher than that in anti-miR-21 or sh-CXCR4 alone group. Next, Flow Cytometry analysis was used to assess the apoptosis in these transfected cells. Statistically more apoptotic cells were found in both anti-miR-21 and sh-CXCR4 groups, and as we expected, anti-miR-21 + sh-CXCR4 group generated the most apoptotic cells (Fig. 3C and Fig. 3D). Thus, our results demonstrated that anti-miR-21 + sh-CXCR4 had an enhanced effect on both suppressing glioma cell proliferation and pro-apoptosis.

**Fig.3.**
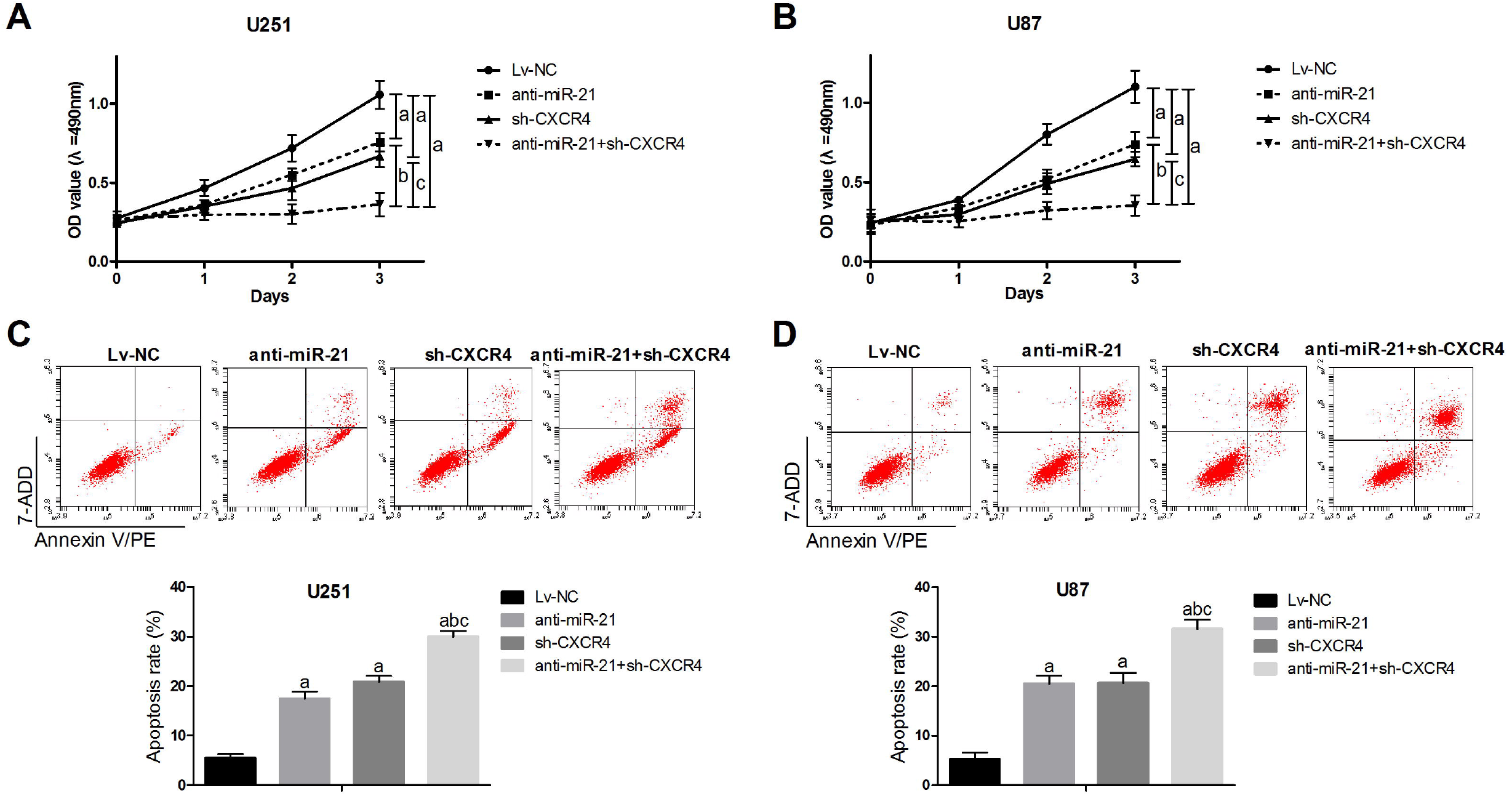
Double targeted knockdown of miR-21 and CXCR4 inhibited proliferation of glioma cells. **(A and B)** MTT assays were employed to assess the proliferation of transfected U251 and U87 cells. **(C and D)** Flow cytometry was used to analyze the apoptosis in transfected U251 and U87 cells. *P* < 0.05, a: *vs* Lv-NC group; b: *vs* anti-miR-21 group; c: *vs* sh-CXCR4 group.

### Double targeted knockdown of miR-21 and CXCR4 inhibited invasion and migration of glioma cells

Transwell assay was performed to evaluate the regulation of miR-21 and CXCR4 on glioma cell invasiveness. Anti-miR-21 or sh-CXCR4 alone was able to significantly limit the invasiveness of glioma cells (Fig. 4A and Fig. 4B). Here our data presented that anti-miR-21 + sh-CXCR4 significantly inhibited the cell invasiveness compared to single-targeted knockdowns (anti-miR-21, and sh-CXCR4) (Fig. 4 A and Fig. 4B). Anti-miR-21 or sh-CXCR4 alone significantly reduced migration in both U87 and U251 cells; while, anti-miR-21 + sh-CXCR4 suppressed inhibited the cell migration compared to single-targeted knockdowns in both cell lines (Fig. 4C and Fig. 4D). This revealed the potent repressive effects of anti-miR-21 + sh-CXCR4 on capability invasion and migration of glioma cells.

**Fig.4.**
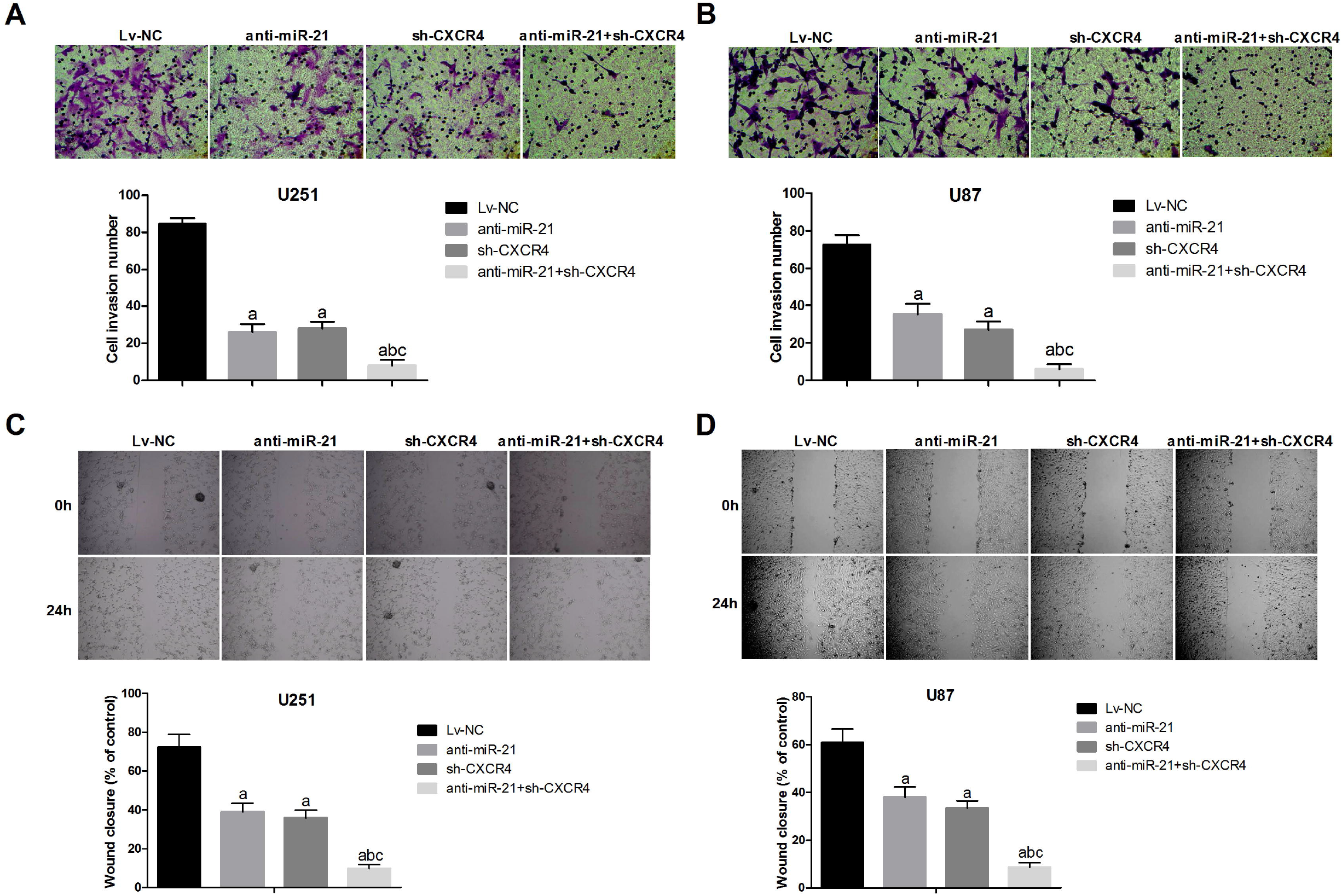
Double targeted knockdown of miR-21 and CXCR4 inhibited invasion and migration of glioma cells. **(A and B)** A Transwell assay was performed on the transfected U251 and U87 cells. Magnification, 200×. **(C and D)** Scratch assays were performed to detect migration of transfected cells. Magnification, 40×. *P* < 0.05, a: *vs* Lv-NC group; b: *vs* anti-miR-21 group; c: *vs* sh-CXCR4 group.

### Double targeted knockdown of miR-21 and CXCR4 slowed tumor growth in glioma xenograft mouse model

To better illustrate the roles of miR-21 and CXCR4 together in tumor growth of glioma, we employed a U87 xenograft mouse model. No obvious difference between the average body weight of mice from different treatment groups (Lv-NC, anti-miR-21, sh-CXCR4, and anti-miR-21 + sh-CXCR4, Fig. 5A). We measured tumor volume every 5 days. The growth of glioma tumor was significantly suppressed in anti-miR-21 or sh-CXCR4 treatment alone groups compared to negative control, while the anti-miR-21 + sh-CXCR4 treatment obviously reduced the tumor growth compared to the anti-miR-21 or sh-CXCR4 group (Fig. 5B and 5C). Furthermore, the miR-21 and CXCR4 level in xenografts were detected. And further, the results had confirmed that the remarkable effect on tumor growth suppression was contributed by inhibition of miR-21 and CXCR4 (Fig. 5D-5F). Thus, our data reported that anti-miR-21 + sh-CXCR4 could diminish growth of glioma xenograft *in vivo.*

**Fig.5.**
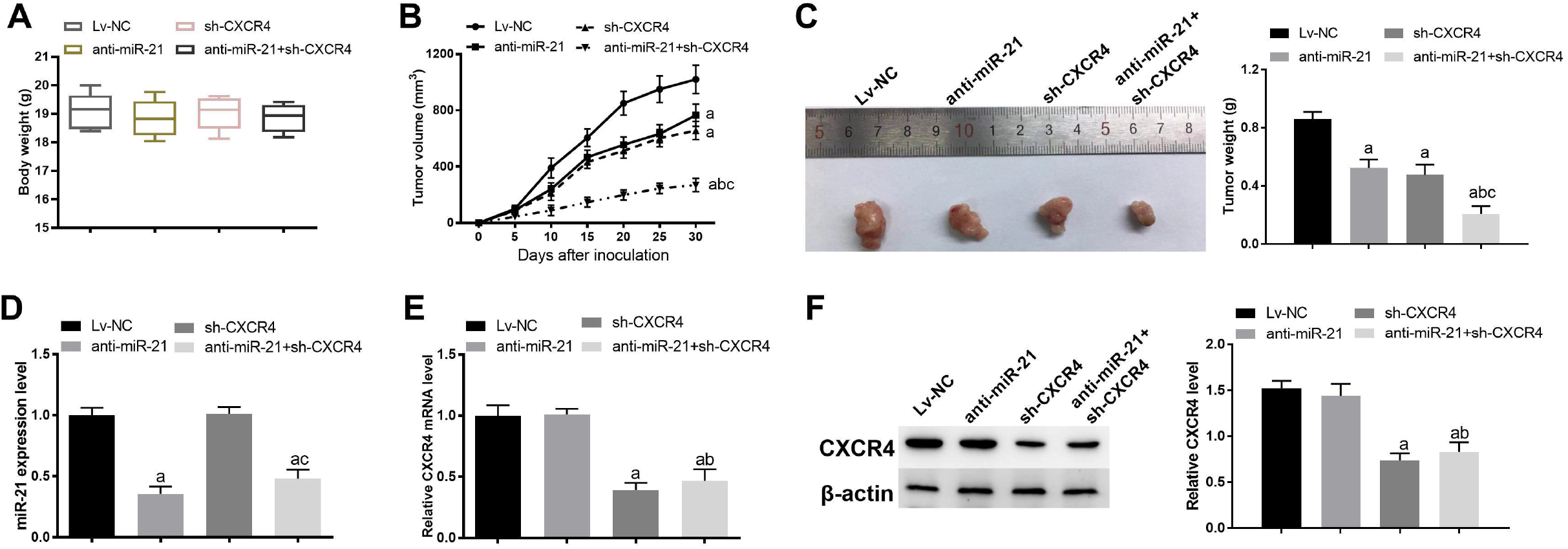
Double-targeted knockdown of miR-21 and CXCR4 inhibited glioma xenograft growth. **(A)** Mice were weighted at 6 weeks post inoculation. **(B)** Tumor volumes were monitored over 4 weeks. **(C)** Tumors were dissected and weighted. **(D and E)** miR-21 and CXCR4 levels in xenograft tumor tissues were measured using qRT-PCR. **(F)** CXCR4 protein levels were tested using western blot. *P* < 0.05, a: *vs* Lv-NC group; b: *vs* anti-miR-21 group; c: *vs* sh-CXCR4 group.

### Double-targeted knockdown of miR-21 and CXCR4 inhibited malignant glioma progression by suppressing of the PI3K/AKT and Raf/ MEK/ ERK pathway

We further investigated the potential downstream pathways that might be responsible for the repressive effect of anti-miR-21 + sh-CXCR4 on tumor progression. In this study, we focused on two pathways 1) PI3K/AKT and 2) Raf/ MEK/ ERK, which both played a vital role in glioma cell fate, and were aberrant in the miR-21 and/or CXCR4 overexpression environment (Ching and Hansel 2010, Guo et al., 2013, Dongfeng et al., 2014, Du et al., 2015, Han et al., 2016, Shao *et al.* 2017). The results from western blot analysis presented no statistically difference in AKT, ERK1/2 among different treatment groups both in glioma cells and xenografts (Fig. 6A-6C). However, anti-miR-21, sh-CXCR4, and anti-miR-21 + sh-CXCR4 all showed a significantly suppressive effect on p-AKT, and p-ERK1/2, with anti-miR-21 + sh-CXCR4 group presenting the strongest inhibition effect in both cell lines and xenografts. These results demonstrated a potent suppressive effect of anti-miR-21 + sh-CXCR4 on activation of AKT axis and ERK axis. These results suggested that double-targeted knockdown of miR-21 and CXCR4 inhibited malignant glioma progression by suppressing of the PI3K/AKT and Raf/ MEK/ ERK pathway.

**Fig.6.**
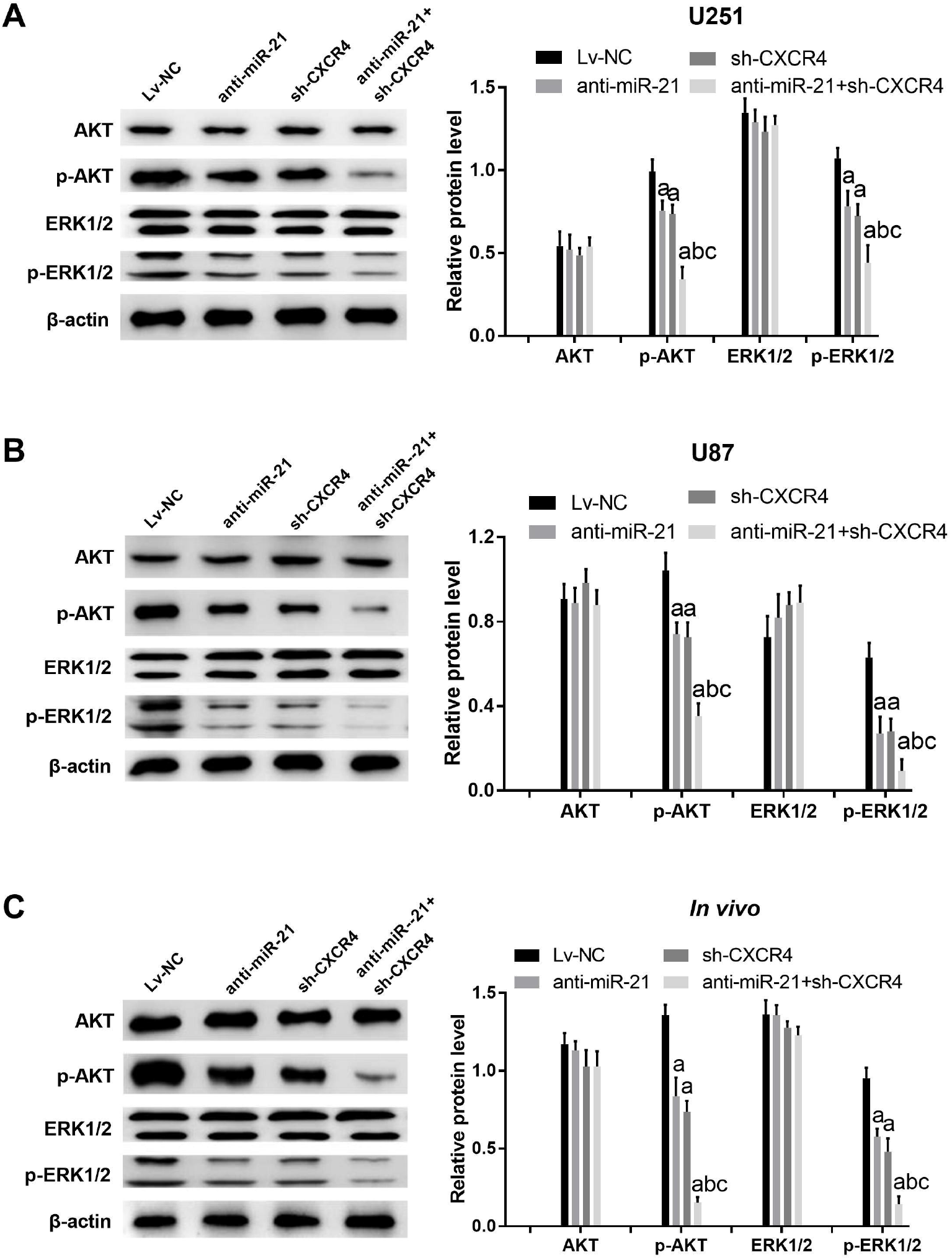
Double-targeted knockdown of miR-21 and CXCR4 inhibited PI3K/AKT and Raf/MEK/ERK pathways in glioma xenograft tissue and cells. **(A-C)** Related proteins were tested using western blot. *P* < 0.05, a: *vs* Lv-NC group; b: *vs* anti-miR-21 group; c: *vs* sh-CXCR4 group.

## Discussion

Glioma is the most common primary malignant brain tumor. Several novel therapies have been developed for the treatment of malignant glioma, including gene therapy, targeted molecular therapy, immunotherapy, and stem therapy. Among them, gene therapy presents great efficiency in suppressing glioma progression (Han *et al.* 2016). Particularly, the combination of multi-gene therapy tends to be more effective than the single-gene one (Jin et al., 2010).

miR-21 is the first miRNA reported involved in human malignant glioma regulation (Melnik 2015). Its expression has been reported positively correlated with clinical grade of glioma malignancy (Lei *et al.* 2014, Hermansen *et al.* 2016). Also, the vital roles of miR-21 in glioma initiation, maintenance, and survival have been proved by numerous studies (Tao *et al.* 2013, Lei *et al.* 2014, Yeh et al., 2015, Hermansen *et al.* 2016, Luo *et al.* 2017, Wang et al., 2017). Studies show that miR-21 downregulation represses cell proliferation and invasiveness, enhances apoptosis, and sensitizes resistant to chemo- and radio-therapy (Loges *et al.* 2012, Lei *et al.* 2014, Wang et al., 2015, Yeh *et al.* 2015, Luo *et al.* 2017, Wang *et al.* 2017). In our study, we confirmed the increased miR-21 in malignant glioma tissues and glioma cells as well as the regulation of it in glioma progression.

CXCR4 is another factor that plays a vital role in glioma pathology (i.e. neoplastic transformation, malignant tumor progression, infiltration, metastasis, angiogenesis, and vasculogeneis) (Dutt et al., 1998, Ono and Freed 1999, Eckert *et al.* 2018). Our data confirmed that CXCR4 expression was up-regulated in malignant glioma tissues and glioma cells in comparison to normal tissue or glial cells. The potential of CXCR4 as a promising therapeutic target for malignant glioma treatment has been extensively investigated and several targeting drugs have already been undergoing clinical trials (Eckert *et al.* 2018). However, most research topics are focused on employing CXCR4 antagonist as a single treatment or in complement with radio- or chemo-therapy (Eckert *et al.* 2018). In our present study, we reported that double-targeted knockdown of miR-21 and CXCR4 presented a better suppressive effect on tumor progression when compared to single-targeted knockdown of miR-21 or CXCR4 alone. Anti-miR-21 + sh-CXCR4 almost abrogated the proliferation, cell invasion, and migration in migration-prone glioma cells. Also, anti-miR-21 + sh-CXCR4 repressed xenograft tumor growth.

Both PI3K/AKT and Raf/MEK/ERK are common and crucial signaling for cancer development (Serena et al., 2011). Activated ATK and ERK phosphorylate and activate downstream proteins involved in regulating cell survival, proliferation and metabolic pathways (Redman et al., 2013, Dueppers et al., 2015). Studies demonstrate that dysregulation of both pathways are observed in various cancers and the inhibition of both pathways would be promising strategy for cancer therapy (Dueppers *et al.* 2015). However, direct inhibition of these pathways is unpractical given the complexity of their crosstalk regulatory mechanisms (Ghayad and Cohen 2010). Interestingly, our results presented that double-targeted knockdown of miR-21 and CXCR4 powerfully suppressed PI3K/AKT and Raf/MEK/ERK activation, which might have significant therapeutic values and offer some insights for further investigation of the molecular mechanism in glioma pathology. Since the overexpression of miR-21 and/ or CXCR4 is also observed in many other cancers, such as, ovarian, gastric, colonic, pancreatic, breast and prostate cancers (Melnik 2015, J Richardson 2016), our study could also shed some light on the efficacious treatment development for them.

## Conclusion

In summary, the knockdown of miR-21 or CXCR4, double-targeted knockdown of miR-21 and CXCR4 could more effectively inhibit the proliferation, migration, invasion and growth of transplanted tumor and promote cell apoptosis, which were involved in the PI3K/AKT and Raf/MEK/ERK signaling pathways. Therefore, double-targeted miR-21 and CXCR4 may be potential and promising therapeutic targets for malignant glioma salvage treatment.

## Material and Methods

### Samples

A total of 25 patients were recruited in this study. All patients were diagnosed with malignant glioma, and diagnosis was confirmed by both clinical symptoms and the histology consensus from at least two neuropathologists at the Department of Oncology in the First Affiliated Hospital of Zhengzhou University. The surgical removed tumor tissues and adjacent peritumoral brain tissues were immediately snap frozen using liquid nitrogen and then stored at −80°C fridge for further qRT-PCR and western blot analysis. Written informed consents for the pathological analyses and experiments were obtained from all of the donors before their surgery. All samples were handled and processed in compliance to the institutional ethical and legal standards, which was approved by the Research Ethics Committee of the First Affiliated Hospital of Zhengzhou University.

### Cell lines and culture

Human glioma cell lines U87 and U251, normal glial cell HAC and embryonic kidney cell HEK-293T were obtained from the American Type Culture Collection (ATCC, USA). All cells were cultured in Dulbecco’s modified Eagle medium (DMEM) /high glucose (Hyclone, Logan, UT) supplemented with 10% FBS (Invitrogen, CarIsbad, CA, USA), 100 U/ml penicillin, and 100 μg/ml streptomycin (Invitrogen) and maintained in humidified incubator at 37 °C with 5% CO_2_ and 95% of air.

### qRT-PCR analysis

Total RNA of cultured glioma cells (U87 and U251) or tissue samples was extracted with the TRIzol reagents (Invitrogen) in accordance with the manufacturer’s instruction. For detecting miR-21 and CXCR4 mRNA, cDNA was first amplified by reverse transcription using the PrimeSCript RT reagent kit (Takara Bio, Inc, Otsu, Japan). Then the relative expressions were measured with TaqMan MicroRNA Assay kit (Applied Biosystems, Foster City, CA, USA) and SYBR-Green PCR Master Mix Kit (Takara) on an ABI7500 system (Applied Biosystem), respectively. The primer sequences used in this study were as followings: CXCR4: sense: 5 ‘-GAAACCCTCAGCGTCTCAGT-3 ‘, antisense: 5’-AGTAGTGGGCTAAGGGCACA-3 ‘; miR-21: sense: 5’-GCGTGTCGGGTAGCTTATCAGAC-3’, antisense from TaqMan MicroRNA Assay kit; GAPDH: sense: 5’-GGGAAACTGTGGCGTGAT-3’, antisense: 5’-GAGTGGGTGTCGCTGTTGA-3’; U6 sense: 5’-CTCGCTTCGGCAGCACATATACT-3’, antisense: 5’-ACGCTTCACGAATTTGCGTGTC-3’. All experiments were performed in triplicate. The relative CXCR4 and miR-21 levels were obtained from comparing to the expression level of GAPDH and *U6* snRNA, respectively. The relative expression levels were calculated using 2^-ΔΔCt^ method.

### Western blot assay

Cultured cell and collected tissue samples were first washed twice with ice-cold PBS and then lysed in RIPA buffer supplemented with protease and phosphatase inhibitor cocktails. 30 μg of protein was separated by 10% SDS-PAGE and then electrophoretically transferred onto a PVDF membrane (Milipore, Bedford, MA). The blots were incubated in 1 × PBS containing 5% skimmed milk powder for 1 hr at room temperature to block the nonspecific binding. Following washing (TBST, 3×, 10 min each), blots were incubated with primary antibodies overnight at 4°C.

After another wash, blots were exposed to horseradish peroxidase-conjugated secondary antibody for 2 hr at room temperature, and visualized using the enhanced chemiluminescence detection kit (Sigma). The following primary antibodies were used in this study: CXCR4 (Abcam, Cambridge, MA, USA), β-actin (Cell Signaling Technology, Danvers, MA, USA), AKT (Cell Signaling Technology), phospho-AKT (Cell Signaling Technology), ERK1/2 (Cell Signaling Technology), and phospho-ERK1/2 (Cell Signaling Technology). The protein levels were digitalized using ImageJ 1.48 version (NIH, Bethesda, Maryland) (Java 1.8.9_66) and relative protein levels were obtained by comparing the protein level to the level of β-actin.

### Construction of the recombinant lentiviral vectors

Negative control and anti-miR-21 sequences were cloned into lentiviral vector (pLent-hU6-EF1-GFP-Puro) to generate the negative control recombinant plasmid (pLenti-Lv-NC) and miR-21 knockdown recombinant plasmid (pLenti-anti-miR-21), respectively. Sh-CXCR4 was cloned into lentiviral vector pLenti-hU6-GFP-Puro (Shanghai sheng-gong) to generate the CXCR4 knockdown recombinant plasmid (pLenti-sh-CXCR4). The double-targeted knockdown plasmid (pLenti-anti-miR-21+sh-CXCR4) was obtained by connected sh-CXCR4 fragment with pLenti-anti-miR-21 plasmid using Ready-to-Use Seamless Cloning Kit (Shanghai sheng-gong).

Before the lentiviral packing, 293 T cells were cultured in 6-well plates until reach 80% confluence. Then, cells were co-transfected with the recombinant lentiviral vectors (Lv-NC, anti-miR-21, sh-CXCR4, and anti-miR-21 + sh-CXCR4) and two auxiliary packaging plasmids (pCMV-Δ8.2 and pCMV-VSV-G)). 72 hours after the transfection, the lentivirus particles were collected by centrifugating at 25000 rpm for 1.5 h at 4°C. All lentiviral constructs were inserted a puromycin resistance sequence for drug screen.

### Lentiviral infection of glioma cells (U87 and U251)

Before the infection, glioma cells (U87 and U251) were routinely cultured and seeded on 6-well plates at a density of 5 x 10^4^ cells/ well. When cell density reached 50% confluence, cells were infected with lentivirus particles (MOI=20) in presence of 8 μg/ml polybrene (GeneCopoeia, Guangzhou, China) for 24 hr. Then the culture were replaced by fresh culture medium containing 2% FBS and cultured for another 72 hr. The effectiveness of lentiviral particles that expressing anti-miR-21, sh-CXCR4, or anti-miR-21 + sh-CXCR4 was examined by qRT-PCR and western blot as described above.

### MTT assay

Logarithmically growing glioma cells (U87 and U251) with different treatments were plated into 96 wells culture clusters (Costar, Cambridge, MA, USA), at a density of 2000 cells/well and cultured for different time points (0, 24, 48, 72 hr). Cell viability was assessed by adding 10 μl MTT (0.5 mg/ml) to each well and incubated for 4 hr. After removing the cell medium, 100 μl DMSO was added and cell proliferation was measured at 490 nm wavelength. Measurements of cell viability for each treatment were done in triplicates from three independent experiments. Treatments included: 1) anti-miR-21, 2) sh-CXCR4, 3) anti-miR-21 + sh-CXCR4, 4) Lv-NC.

### Flow cytometry analysis

Annexin V-PE/7-AAD Apoptosis detection Kit I (BD Bioscience, USA) was employed to determine the apoptosis of transfected glioma cells (U87 and U251). Cells were harvested after transfected with lentivirus, washed twice with PBS, and then resuspended in 400 μl binding buffer. After the resuspension, cells were incubated with 5 μl Annexin V-PE and 10 μl 7-AAD solutions at room temprature for 10 min in the dark. The analysis was performed by flow cytometry (FACSCalibur; Becton Dickinson). For each sample, 10,000 cells were analyzed. All of the experiments were performed in triplicate. Transfection treatments include: 1) anti-miR-21, 2) sh-CXCR4, 3) anti-miR-21 + sh-CXCR4, 4) Lv-NC.

### Transwell assay

The invasion of glioma cells (U87 and U251) was assayed using BioCoat Matrigel invasion chambers (BD Bioscience) with poly carbonic membrane (6.5 mm in diameter, 8 μM pore size). Before the start of the assay, the chambers were inserted into 24-well culture plates and pre-treated with serum-free DMEM medium at 37°C for 30 min. In the Transwell assay, the transfected cells were resuspended in 300 μl serum-free DMEM at a density of 5 × 10^4^ cells/ml and added to the upper chamber. 500 μl DMEM supplemented with 20% FBS was added into the lower chamber of each well. The plates were incubated in humidified incubator (37°C, 5% CO_2_) for 20-24 hr. After the incubation, non-migrated cells in the upper chamber were removed mechanically with a cotton swap, while the invaded cells in the lower side of the membranes were fixed with 5% glutaraldehyde for 15 min at 4 °C, washed with PBS twice, and stained with 1% crystal violet for 15 min. Pictures were photographed at 200× magnification, and cell numbers were counted from at least five randomly selected fields.

### Scratch assay

Transfected glioma cells (U87 and U251) were suspended and plated in 6-well plates until they reach 70-80% confluence. At the initial time, a scratch was made on the cell monolayer using a sterile 200 μl micropipette tip. Pictures were photographed at different time points (0, 24 hr), with 40 × magnification.

### Generation of tumor xenografts and in vivo treatments

All the animal procedures were approved by Ethics of Animal Experiments of the First Affiliate Hospital of Zhengzhou University, and all the experiments were performed in strict accordance with the institutional guidelines. BALB/C nude mice weighting 21.2±3 g and being 5 weeks old were housed in sterile cages under standard conditions (23 ± 2°C, 45-55% humidity, 12 hr light duration). The mice were allowed for free access to water and acclimatized for one week before the experiments. The mice were randomly divided into 4 groups. Immediately prior to the xenograft experiment, the transfected (Lv-NC, anti-miR-21, sh-CXCR4, and anti-miR-21+sh-CXCR4) cells were suspended to single-cell in sterile saline. The transfected U87 cells were injected into the left axillary subcutaneous location of BALB/C nude mice. The xenograft procedure was designed according to the published ones (Qin et al., 2017, Li et al., 2018). In brief, immediately prior to the xenograft experiment, the transfected cells were suspended to single-cell in sterile saline. Following the surgery, mice were monitored for tumor growth every 5 days. The formula V (volume) = (long diameter × short diameter^2)/2 was used to calculate the tumor volume. Mice were weighted and sacrificed at 4 weeks post-xenograft; then tumors were collected and measured. Immediately after the measuring, xenografts were stored at −80°C for further qRT-PCR and western blot analysis.

### Statistical analysis

Statistical analysis was performed with SPSS 19.0 with an overall significance level of *P* < 0.05. Differences between the groups were computed by two-tailed unpaired Student t-Test.

## Acknowledgements

No.

## Conflict of interest

No potential conflicts of interest were disclosed.

